# Morphological simulation tests the limits on phenotype discovery in 3D image analysis

**DOI:** 10.1101/2024.06.30.601430

**Authors:** Rachel A. Roston, Sophie M. Whikehart, Sara M. Rolfe, Murat Maga

## Abstract

In the past few decades, advances in 3D imaging have created new opportunities for reverse genetic screens. Rapidly growing datasets of 3D images of genetic knockouts require high-throughput, automated computational approaches for identifying and characterizing new phenotypes. However, exploratory, discovery-oriented image analysis pipelines used to discover these phenotypes can be difficult to validate because, by their nature, the expected outcome is not known *a priori*. Introducing *known* morphological variation through simulation can help distinguish between real phenotypic differences and random variation; elucidate the effects of sample size; and test the sensitivity and reproducibility of morphometric analyses. Here we present a novel approach for 3D morphological simulation that uses open-source, open-access tools available in 3D Slicer, SlicerMorph, and Advanced Normalization Tools in R (ANTsR). While we focus on diffusible-iodine contrast-enhanced micro-CT (diceCT) images, this approach can be used on any volumetric image. We then use our simulated datasets to test whether tensor-based morphometry (TBM) can recover our introduced differences; to test how effect size and sample size affect detectability; and to determine the reproducibility of our results. In our approach to morphological simulation, we first generate a simulated deformation based on a reference image and then propagate this deformation to subjects using inverse transforms obtained from the registration of subjects to the reference. This produces a new dataset with a shifted population mean while retaining individual variability because each sample deforms more or less based on how different or similar it is from the reference. TBM is a widely-used technique that statistically compares local volume differences associated with local deformations. Our results showed that TBM recovered our introduced morphological differences, but that detectability was dependent on the effect size, the sample size, and the region of interest (ROI) included in the analysis. Detectability of subtle phenotypes can be improved both by increasing the sample size and by limiting analyses to specific body regions. However, it is not always feasible to increase sample sizes in screens of essential genes. Therefore, methodical use of ROIs is a promising way to increase the power of TBM to detect subtle phenotypes. Generating known morphological variation through simulation has broad applicability in developmental, evolutionary, and biomedical morphometrics and is a useful way to distinguish between a failure to detect morphological difference and a true lack of morphological difference. Morphological simulation can also be applied to AI-based supervised learning to augment datasets and overcome dataset limitations.

## 1 Introduction

International efforts to characterize mammalian genes and phenotypes have rapidly grown the amount of 3D imaging data available for phenotype discovery in knockout mouse lines (Dickinson *et al*., 2016; Groza *et al*., 2023). As of Data Release 21.0, the International Mouse Phenotyping Consortium (IMPC) has phenotyped over 8900 genes and 9500 mutant mouse lines (www.mousephenotype.org). Reverse genetic screens, such as those conducted by the IMPC, have traditionally focused on identifying overt morphological differences. But, advances in 3D imaging, such as the adoption of diffusible-iodine contrast-enhanced micro-CT scanning (diceCT) by the IMPC, allow new opportunities for quantitative image analysis and high-throughput approaches for phenotype discovery (Nieman, Wong and Henkelman, 2011; Norris *et al*., 2013; Wong *et al*., 2014; Dickinson *et al*., 2016; Horner *et al*., 2021; Handschuh and Glösmann, 2022). Quantitative computational approaches support the identification of more subtle phenotypes often missed by qualitative observation; the detection of novel phenotypes at earlier developmental stages, before they become apparent to human observers; and systematization and automation. 3D image analysis techniques for phenotype discovery may include automated segmentation of organs to calculate their volumes (*e.g.*, Wong *et al*., 2014; Rolfe, Whikehart and Maga, 2023), comparing size and shape of regions of interest using geometric morphometrics (*e.g.*, Devine *et al*., 2022), comparing voxel intensities (*e.g.*, Ashburner & Friston 2000; Wong *et al*., 2014), or comparing deformation fields and their derivatives (*e.g.*, Horner *et al*., 2021; Roy *et al.,* 2013).

Tensor-based morphometry (TBM) is a statistical methodology to calculate differences in tissue volume of two groups from 3D images. This is achieved by deformably registering a series of images to a reference (*i.e.*, template) that forms the statistical and geometric basis of comparison, and then calculating Jacobian determinants of the resultant vector fields for each subject (Wong *et al*., 2014; Horner *et al*., 2021). A statistical model applied to the Jacobian determinants indicates the relative volumetric change (shrinkage and expansion) and its magnitude in a per-voxel basis using the reference image. Since the significance of the difference is evaluated on a per-voxel basis, a multiple-test correction penalty must be applied to the results. TBM has been used to study a variety of research questions such as the identification of differences in brain morphology in fetal alcohol spectrum disorder, HIV/AIDS, and childhood-onset schizophrenia (Chiang *et al*., 2007; Gogtay *et al*., 2008; Meintjes *et al*., 2014) and differences in embryonic size and morphology in mice (Zamyadi *et al*., 2010; Horner *et al*., 2021).

Using TBM for exploratory analysis of uncharacterized phenotypes can be challenging because expected outcomes are not known *a priori.* A failure to detect differences may truly reflect a lack of morphological difference, but it may also result from the relatively high penalty for multiple comparisons, low sample sizes, a low effect size, or a combination of these issues. Distinguishing between these scenarios is an essential step in validating image analysis pipelines. To empirically test pipelines, researchers usually need to find a mutant line with a well-characterized phenotype and hope that their image analysis pipeline identifies the same known phenotype (Wong *et al*., 2014). If known phenotypes are recovered, researchers can apply the analysis pipeline to mutant lines that were not previously characterized. While this sort of empirical validation is useful, it cannot directly address the issues we have mentioned above. And, more importantly, it limits the validation of image analysis pipelines to a few well-characterized mutant lines that may not exhibit all possible anatomical variations and phenotypes which can be independently identified and characterized.

An alternative, far more flexible approach to validation of image analysis pipelines is to create datasets with a “known signal”—*i.e*., simulated anatomical differences that vary in their effect size—and then analyze these simulated datasets to empirically investigate issues around sample size, effect size, and multiple-comparison penalty. Morphological simulation can help distinguish whether results reflect a computational artifact or real morphologic differences and can also test the sensitivity and reproducibility of analyses. Previous approaches to simulation have focused primarily on parametric models of tissue atrophy and growth (*e.g.*, Karaçali and Davatzikos, 2006; Rolfe *et al*., 2011). However, these approaches do not allow simulations of more complex morphological differences that differentially affect multiple body regions.

Here we describe a fairly simple methodology for generating 3D simulated morphological differences that allows researchers to directly shape the types of local variations they wish to introduce. Using open-access data and open-source tools available in 3D Slicer, SlicerMorph, and Advanced Normalization Tools in R (ANTsR) makes this approach broadly accessible and reproducible (Fedorov *et al*., 2012; Kikinis, Pieper and Voxburgh, 2014; Rolfe *et al*., 2021; Tustison *et al*., 2021). In the present paper, we primarily focused on creating quantitative differences in a single organ (the heart). We focus on quantitative differences because gross structural defects are easy to identify through conventional necropsies under the microscope, whereas quantitative differences in size and shape of an organ require careful measurements, which are not feasible without this type of 3D imaging. We then use the original and simulated data to test the conditions under which we can reliably detect induced differences using TBM. Our results show that our method of simulation retains individual variability and that TBM is able to recover introduced differences under certain conditions.

## 2 Materials and Methods

### 2.1 Materials

Thirty wildtype (“baseline”) diceCT scans of E15.5 mouse embryos were randomly selected from the International Mouse Phenotyping Consortium (IMPC) database (www.mousephenotype.org). A 3D reference image (*i.e.*, template) of E15.5 embryos derived from diceCT scans of wildtype mice, along with its atlas with 50 annotated anatomical structures, was previously published by Wong et al. (2012). We utilize this template as the basis of our statistical analysis and, along with its corresponding atlas, to simulate differences in heart volume and rostrum length.

### 2.2 Generating Simulated Deformations

Simulated morphological deformations were created in the following manner: an E15.5 diceCT scan (the subject) was deformably registered to the E15.5 template using Advanced Normalization Tools in R (ANTsR) (Avants, Tustison and Song, 2009; Tustison *et al*., 2021). This registration provided a mapping of subjects into the template’s reference frame. While in the template’s reference frame, we applied a simulated deformation that we had created to the subject, and then propagated the deformed subject back to the subject space by applying the inverse transform of the deformable registration.

To generate the simulated deformations in the template space, we first created a model of the external body surface from the template and a model of the surface of the heart ventricles from the E15.5 template and atlas in 3D Slicer using the available segmentation tools (Fedorov *et al*., 2012; Kikinis, Pieper and Voxburgh, 2014). Then, on these models, we sampled uniformly spaced pseudolandmarks (Figure 1A,B) using the PseudoLMGenerator module from the SlicerMorph extension (Rolfe, Davis and Maga, 2021; Rolfe *et al*., 2021). A copy of the original heart pseudolandmark set was scaled according to simulation experiments in Table 1. Unmodified external body surface pseudolandmarks and unmodified heart pseudolandmarks were then merged to create a composite unmodified pseudolandmark set (Figure 1C). Each scaled heart pseudolandmark set was also merged with a copy of the unmodified external body surface pseudolandmarks to create modified pseudolandmark sets (Figure 1C). To simulate rostrum elongation, the external body surface pseudolandmark set was cloned and landmarks at the end of the rostrum were moved distally, increasing the length of the rostrum by approximately 20%. Simulated deformation transforms were then generated by registering the unmodified pseudolandmark set to each modified pseudolandmark set (Figure 1D).

**Figure 1.**
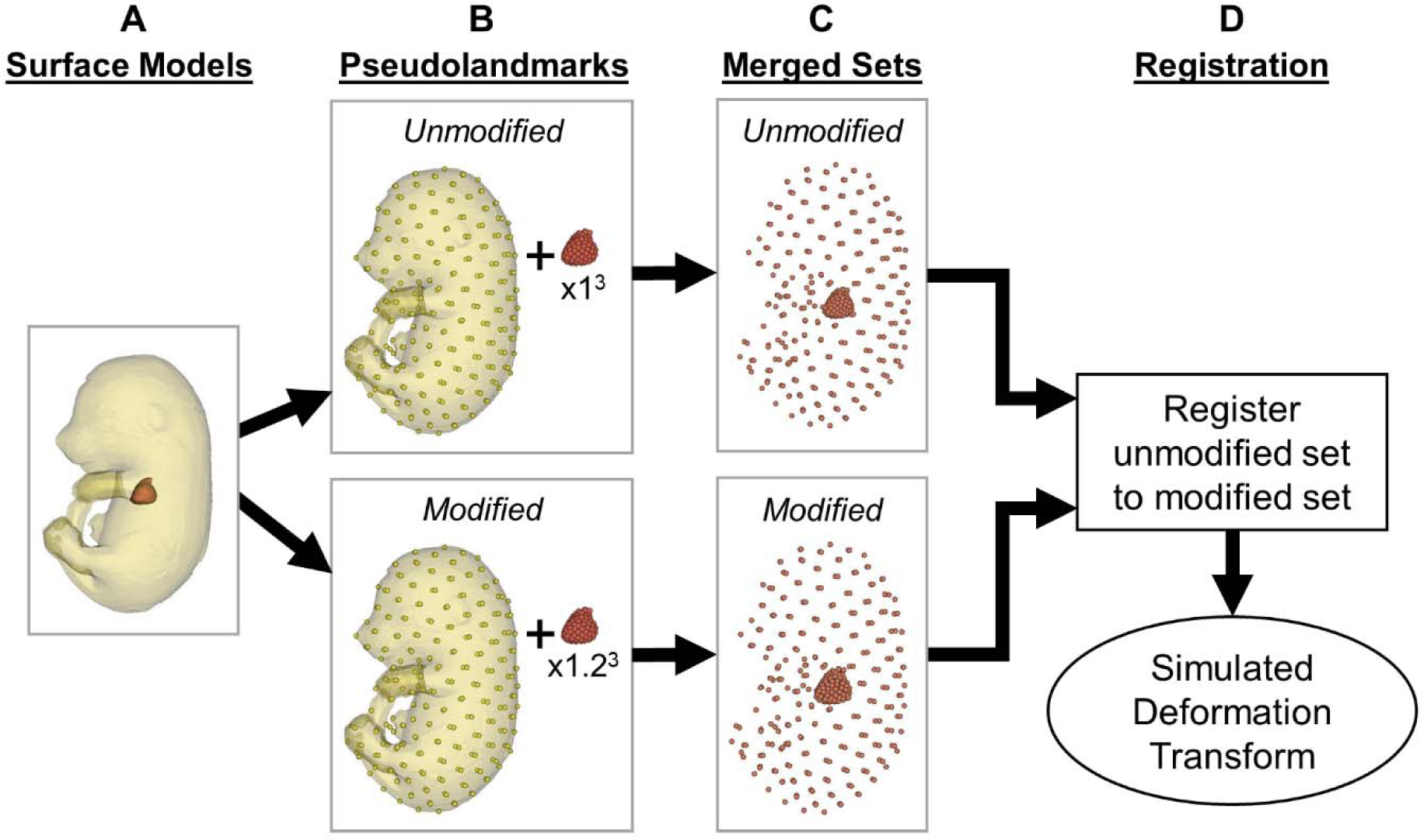
Generation of the simulated deformations, demonstrated with heart expansion with a scale factor of 1.2^3^ (to 173% of its original volume) as an example.

**Table 1.** Simulated deformations included in this study.

| **Body Part** | **pseudoLM Sets** | **Simulated Deformation** | **Scale Factor** | **Expected Percentage of Original Volume** |
| --- | --- | --- | --- | --- |
| Heart Ventricles | External body, heart ventricles | Volume Reduction | $0.80^3$ | 51% |
| Heart Ventricles | External body, heart ventricles | Volume Reduction | $0.90^3$ | 73% |
| Heart Ventricles | External body, heart ventricles | Volume Reduction | $0.95^3$ | 85% |
| Heart Ventricles | External body, heart ventricles | Volume Expansion | $1.05^3$ | 116% |
| Heart Ventricles | External body, heart ventricles | Volume Expansion | $1.10^3$ | 133% |
| Heart Ventricles | External body, heart ventricles | Volume Expansion | $1.20^3$ | 173% |
| Rostrum | External body | Elongation | $\sim 1.20$ | $\sim 120\%$ length increase |

### 2.3 Volumetric Analysis

Control (*i.e.,* original) and experimental (simulation) subjects were segmented using the MEMOS module in 3D Slicer (Rolfe, Whikehart and Maga, 2023). Segment volumes were calculated using the labelStats function of ANTsR and plotted with ggplot2 in R (Avants, Tustison and Song, 2009; Wickham, 2016; RStudio Team, 2020; Tustison *et al*., 2021; R Core Team, 2023).

### 2.4 Tensor Based Morphometry (TBM)

Control (*i.e.*, original) and experimental (*i.e.*, simulated) subjects were deformably registered to the E15.5 template and Jacobian determinants were calculated from the deformable transform for each subject using the CreateJacobianDeterminantImage function of ANTsR (Avants, Tustison and Song, 2009; Wong *et al*., 2012; Tustison *et al*., 2021). A voxel-wise linear regression was performed and p-values were adjusted for multiple comparisons using false discovery rate (FDR, 0.05) by using the relevant functions in the ANTsR and base R libraries (Avants, Tustison and Song, 2009; Tustison *et al*., 2021; R Core Team, 2023). We used two different masks to define two regions of interest (ROIs): the whole body and the thoracic cavity.

## 3 Results

### 3.1 Volumetric Analysis

Heart volume reduction and expansion were each simulated at three magnitudes: 51%, 73%, 85%, 116%, 133%, and 173% of the original heart volume of each baseline subject. Original images with no simulated deformation are denoted as 100% of their original volume. These simulations were confirmed by calculating heart volumes from segments generated from MEMOS (Figure 2). At the population level, simulations of heart expansion and reduction produced the expected changes in volume. However, the mean magnitude of reduction and expansion measured from MEMOS segments was slightly different than the magnitude that we simulated. For instance, simulating a reduction of heart volume to 51% of its original size produced a measured reduction to ∼57% of the heart’s original size. And, simulating an expansion of heart volume to 173% of its original size produced a measured expansion to ∼155% of its original size. The precise amount of reduction and expansion varied slightly in each subject and subjects retained their individual variability (Figure 2). The distributions of heart volumes were minimally affected (Figure 2).

**Figure 2.**
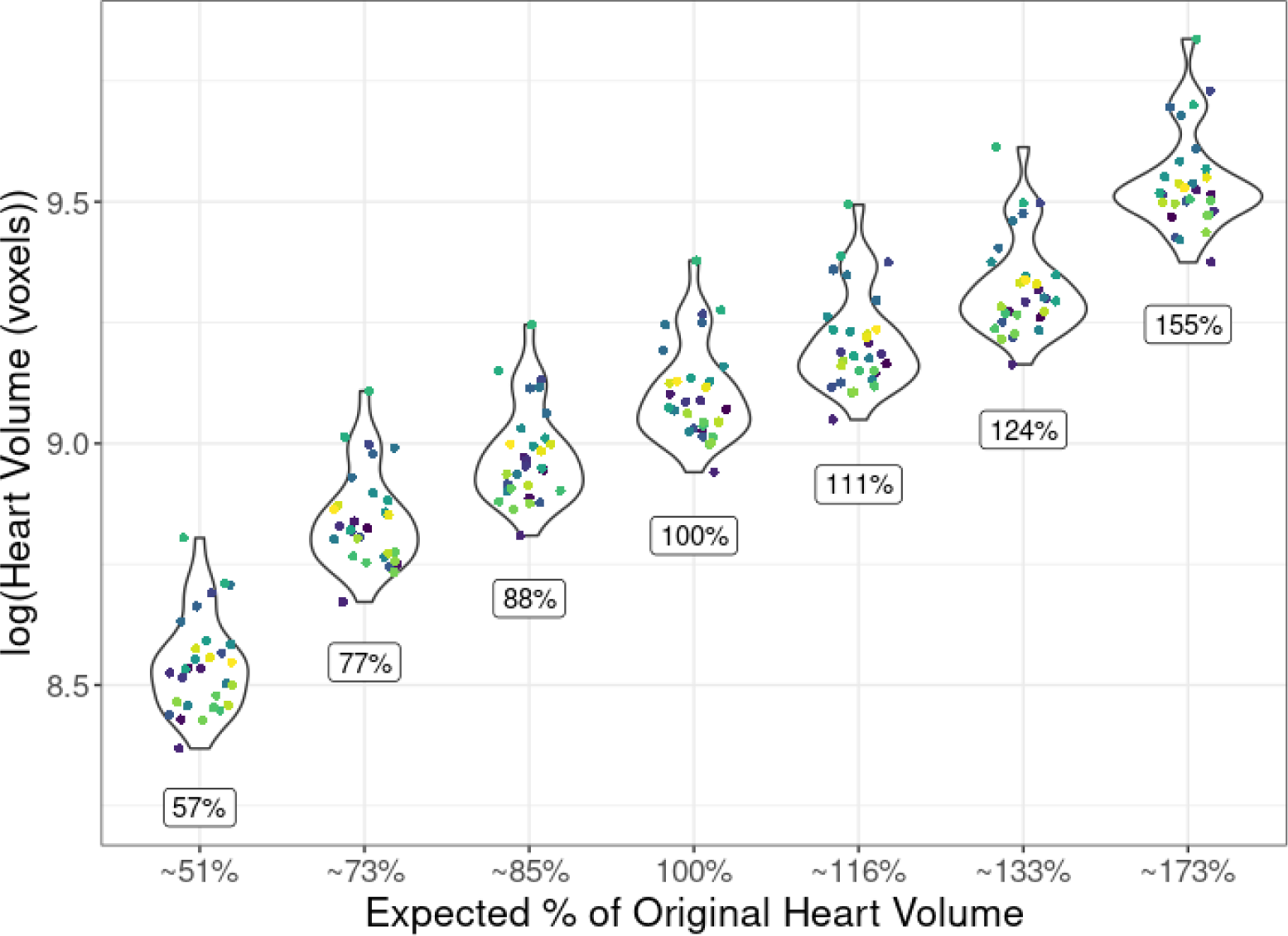
Experimental heart volumes measured from MEMOS segmentation. Mean heart volumes (indicated by boxed numbers) increased and decreased as expected with simulation of expansion and shrinking. Population-level variability in heart volume was retained.

### 3.2 Tensor-Based Morphometry (TBM)

#### 3.2.1 Heart Volume Reduction and Expansion

The TBM pipeline detected volumetric changes in two of the heart reduction (51% and 73%) and two of the heart expansion (133% and 173%) experiments (Table 2). Simulations with a small magnitude of reduction or expansion (*i.e.*, 85% and 116%) were not detected with our analytical pipeline even when all thirty experimental images were included in the analysis.

**Table 2.** Minimum sample size required for consistent detection of heart volume simulations through tensor-based morphometry (TBM). Experimental sample sizes of n=3, 6, 9, 12, 15, 18, 21, 24, 27, and 30 were compared with thirty original baseline subjects. For each sample size, five sets of subjects were generated through random sampling and overlap between sets was minimized. The minimum sample size for detection in all five sets of subjects is reported.

| Simulated Deformation | Expected Percentage of Original Volume | Region of Interest |  |
| --- | --- | --- | --- |
|  |  | Whole Body | Thorax |
| Reduction | 51% | 6 | 3 |
| Reduction | 73% | 27 | 12 |
| Reduction | 85% | Not detected | Not detected |
| Expansion | 116% | Not detected | Not detected |
| Expansion | 133% | 27 | 18 |
| Expansion | 173% | 6 | 3 |

When the ROI for analysis was set to the entire body, the largest magnitudes of simulated reduction and expansion could be detected with sample sizes of only six subjects (Table 2). As expected, when we simulated smaller volumetric changes, a larger sample size of twenty-seven subjects was required to detect differences (Table 2). Setting the ROI to the thorax only, rather than the whole body, lowered the minimum sample size required for detection by nearly half for each magnitude of simulated volume change.

Each TBM analysis was replicated at least five times using different sets of subjects in each replicate. For each sample, heatmaps showed significantly different (p < 0.05) volumetric changes in the heart region, as expected, but the precise locations of significantly different voxels within the heart region varied slightly depending on which subjects were included in the sample (Figure 3). Nevertheless, the minimum sample size required for detection was highly consistent, regardless of which subjects were included in the sample (data not shown; for additional information see Roston *et al*., 2024).

**Figure 3.**
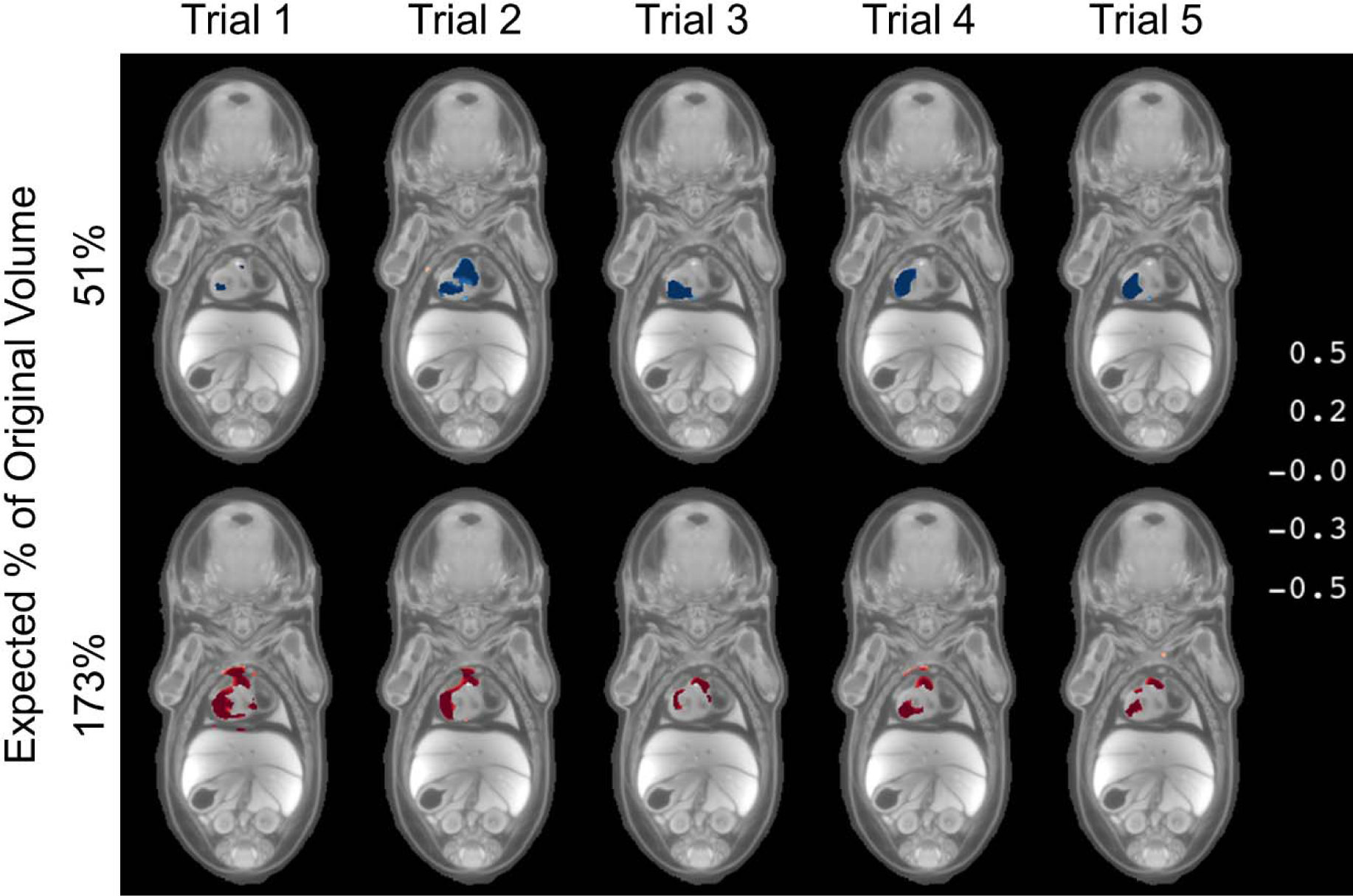
Reproducibility of TBM detection of simulated deformations with a whole-body region of interest. TBM analyses were performed on five non-overlapping sets of six randomly selected subjects. Results consistently showed significantly different (FDR, 0.05) Jacobian determinants in and around the heart, but the precise location of significantly different Jacobian determinants varied based on which six subjects were included in the sample.

#### 3.2.2 Elongation of the Rostrum

To confirm that morphological simulations at the edge of the body can also be detected with TBM, we also simulated elongation of the rostrum to ∼120% of its original length (Figure 4A). TBM detected this elongation with a sample size of three subjects (replicated 10 times; see Roston *et al*., 2024). Heatmaps showed the strongest signal when the sample size was six or greater (Figure 4).

**Figure 4.**
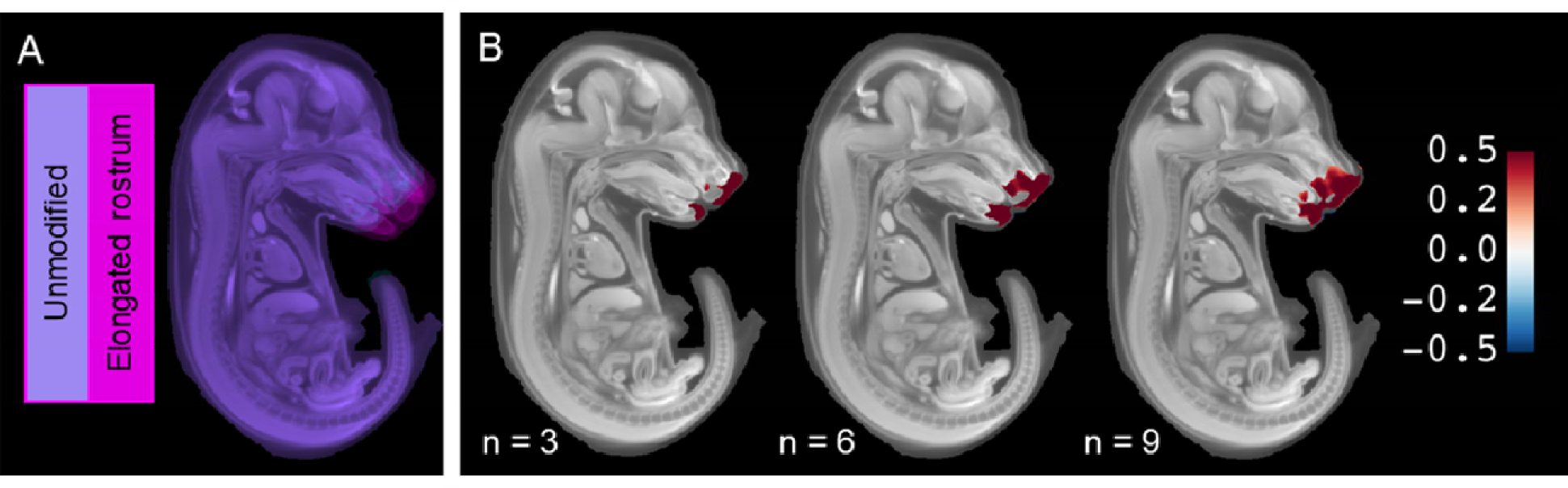
TBM detected simulated elongation of the rostrum to ∼120% its original length. (A) Reference image (template) with simulated deformation of an elongated rostrum (magenta) overlaid over the unmodified reference image (cyan). The lack of cyan voxels in the image indicates that only the rostrum has been modified by the simulation. (B) TBM consistently detected elongation of the rostrum with a sample size of three or more subjects, but the signal increased as more subjects were added to the analysis.

## 4 Discussion

### 4.1 A novel method for morphological simulation

To benefit from the rapidly growing abundance of micro-CT data, high-throughput and automated approaches for phenotype discovery are required (Zamyadi *et al*., 2010; Roy *et al*., 2013; Wong *et al*., 2014; Dickinson *et al*., 2016; Horner *et al*., 2021). However, validating the results of automated pipelines for phenotype discovery can be challenging. Quantitative phenotypes detected by computational pipelines are likely not directly observable to the human eye. Sample sizes for knockout lines of key developmental genes are often small because the mutations are lethal. Hence, real phenotypic differences may go undetected because statistical power is insufficient. Effect sizes may be small, especially when phenotypes are incompletely penetrant and quantitative (Horner *et al*., 2021). Voxel-based image analysis using the entire subject as the analytical ROI may carry a large penalty for multiple comparisons. The typical approach for validating computational pipelines uses well-characterized mutants (*e.g.*, Wong *et al*., 2014), but these well-characterized phenotypes may not be sufficiently diverse to assess the efficacy of a pipeline for all possible differences that might be induced through genetic mutation.

Morphological simulations allow the introduction of known morphological differences with known directionality and effect size, providing *a priori* knowledge of what differences an analytical pipeline for phenotype discovery should detect. Thus, they can be a powerful tool for validating automated phenotyping pipelines and for investigating the limitations of image analyses. Previous approaches to morphological simulation have focused on parametric approaches, limiting simulated morphologies to those which can be generated mathematically (*e.g.*, Karaçali & Davatzikos 2006; Rolfe et al. 2011). Our novel approach instead allows researchers to directly manipulate landmarks to generate simulated deformations that are biologically-informed and which mimic realistic phenotypes that are anticipated in a dataset. These simulated deformations are easy and intuitive to generate from a reference image using the open-access, open-source tools in 3D Slicer, SlicerMorph, and Advanced Normalization Tools for R (ANTsR) (Fedorov *et al*., 2012; Kikinis, Pieper and Voxburgh, 2014; Rolfe *et al*., 2021; Tustison *et al*., 2021). Once generated from the reference image, simulated deformations are propagated to individual subjects using inverse transforms generated from image registration of the subjects to the reference. Although the same deformation is applied to all subjects while they are in template space, the final outcome of deformation for each subject is different because it depends on how similar or different the subject is to the reference image. The variability of simulated deformation is clearly visible in Figure 2; if there were no individual variability in the outcome of heart volume modification, all points within a simulated experiment would have followed a perfect correlation between the expected and the measured heart volume. Thus, our results show that these simulated morphological datasets mimic real-life morphological datasets because they retain individual variability and population-level distributions. Furthermore, our results confirm that simulated expansion and reduction can be detected with volumetric analysis and tensor-based morphometry (TBM) (Table 2; Figures 3, 4).

### 4.2 Detecting Phenotypic Differences with TBM

TBM can be a powerful technique for detecting regional volumetric differences in 3D images (Chiang *et al*., 2007; Gogtay *et al*., 2008; Zamyadi *et al*., 2010; Meintjes *et al*., 2014; Wong *et al*., 2014; Horner *et al*., 2021). Through simulation, we found that the detection of phenotypes through TBM is highly dependent on (i) the magnitude of the phenotypic difference (effect size), (ii) the number of subjects included the analysis (sample size), and (iii) the ROIs included in the analysis, which affects the multiple-comparison penalty. As expected, detection improved with larger sample sizes. But, especially in genetic screens of developmentally essential genes, generating large sample sizes to detect small effect sizes may not be feasible. Our simulations also showed that limiting the number of voxels included in the analysis through the use of ROIs can substantially reduce the sample size required for detection of subtle phenotypes. This demonstrates that the detectability of subtle phenotypes can be substantially improved through a methodical use of ROIs, particularly when there is biological insight that the effect is likely on a specific anatomical system, rather than being on the whole subject.

Our simulated datasets also allowed us to test the reproducibility of our TBM results (Figure 3). When we simulated heart reduction, heatmaps variably highlighted voxels toward the center of the heart, and when we simulated heart volume expansion, heatmaps variably highlighted voxels at the periphery of the heart (Figure 3). This shows that regions of statistical significance do not always occur directly in the regions affected by the simulation and that detected regions of difference can vary somewhat based on which subjects were included in an analysis (Figure 3). Thus, the most appropriate use of TBM is to identify the *general region* of volumetric differences rather than specific voxels. In other words, differences identified by TBM are a hypothesis that should be confirmed through other independent analyses.

### 4.3 Simulated morphology and the future of phenotype discovery

As we have demonstrated, morphological simulation is a useful tool for testing the sensitivity, reproducibility, and limitations of high-throughput, automated phenotype discovery pipelines. Gross morphological differences are easy to identify visually. Therefore, in this study, we focused on introducing subtle, volumetric differences of a type most likely to be targeted by quantitative analyses and most difficult to detect with an automated phenotype discovery pipeline. These simulations were relatively conservative, altering only one organ or facial region in a manner that supports successful image registration. The success of image registration directly influences the ability of TBM to detect meaningful morphological differences (Wong *et al*., 2014; Horner *et al*., 2021). Fully testing the limits of TBM and other phenotype discovery pipelines will require creative introduction of novel morphologies through simulation.

Our approach for morphological simulation—altering landmark positions to generate simulated deformations—is highly adaptable to introduce known differences other than differences in volume, including differences in shape, topology, and angles of different body regions. For this reason, it has great potential to aid in the continued development of image analysis methods to rapidly screen for non-volumetric differences. These may include differences in angle or curvature and differences in developmental rates.

Morphological simulation through the generation of simulated deformations also has applications for the augmentation of datasets for deep learning. Data augmentation is an important method widely used to improve the accuracy and robustness of deep learning models by making changes to the data that are representative of the differences that are expected to be observed between subjects that are not in the training dataset (*e.g.*, Hussain *et al*., 2017; Tustison *et al*., 2019; Nanni *et al*., 2021). Augmentation often involves applying simple transforms such as changes in voxel intensity, cropping, rotations, and resizing (*e.g.*, Hussain *et al*., 2017; Nanni *et al*., 2021; Mulqueeney *et al*., 2024). Additionally, methods for representing more complex shape differences based on the training data have been implemented using transforms derived from PCA of the modes of shape change in a dataset (*e.g.*, Nanni *et al*., 2021). However, all of these augmentation methods still depend on whether desired shape changes already exist in the training data. The methods presented in the present manuscript provide a route to create realistic deformations of anticipated anatomical differences that may not be represented in training data or real biological datasets.

### 4.4 Summary

Validating image analysis pipelines is an essential step in developing new approaches for high-throughput, automated phenotype discovery. Here, we present a novel approach for generating image data with known morphological differences through morphological simulation. Our approach uses open-access, open-source tools available in 3D Slicer, SlicerMorph, and ANTsR to generate a simulated deformation from landmarks and propagate the simulated deformation to each individual subject. This generates realistic simulated datasets by shifting the average morphology of the population in a known manner while retaining individual variability. Simulated morphological data can then be used to test the sensitivity and reproducibility of phenotype discovery pipelines. Here, we demonstrated the use of simulated data to investigate how effect size, sample size, and ROIs affect the sensitivity of TBM to detect introduced morphological differences. The results demonstrated that methodical use of ROIs can greatly enhance the ability of TBM to detect subtle differences. We also tested the reproducibility of TBM results when different sets of individuals and sample sizes were used in the analysis. While the precise location of voxels identified varied between replicates, TBM consistently detected the introduced morphological differences. These results indicate that TBM is a useful technique for identifying general regions of difference, but not necessarily precise locations of difference. In addition to testing and validating image analysis pipelines for phenotype discovery as demonstrated here, morphological simulation can also be used to augment morphological datasets for deep learning. Overall, this makes morphological simulation an essential tool in the continued development of approaches for computational anatomy.

## Acknowledgments

We thank C. Zhang, O. Thomas, M. Bell, and D. Mao for valuable discussion while preparing this manuscript.

## Author Contributions

RAR conceptualized and designed the study, generated simulations, analyzed and interpreted data, and drafted/revised the manuscript. SMW contributed data for the manuscript. SMR conceptualized and designed the study, contributed code for simulations and dataanalysis, and drafted/revised the manuscript. AMM conceptualized and designed the study, contributed code for simulations and data analysis, and drafted/revised the manuscript.

## Funding

This research was partly funded by grants from the National Science Foundation (DBI-2301405) and National Institutes of Health (NICHD-HD104435).

## Data Availability

All 3D data used in the study is publicly available from the data portal of the International Mouse Phenotyping Consortium (https://www.mousephenotype.org). The E15.5 baseline mouse population average reference image was acquired from Toronto Center for Phenogenomics https://www.mouseimaging.ca/technologies/mouse_atlas/mouse_embryo_atlas.html). The analytical code and landmark sets to generate the simulated deformations and for the statistical analysis can be found at https://github.com/raroston/SimMorph.

## Notes

https://github.com/raroston/SimMorph

### Competing Interest Statement

The authors have declared no competing interest.

